# Neural representations of others’ traits predict social decisions

**DOI:** 10.1101/2021.08.01.454671

**Authors:** Kenji Kobayashi, Joseph W. Kable, Ming Hsu, Adrianna C. Jenkins

**Affiliations:** University of Pennsylvania; University of California, Berkeley

## Abstract

To guide social interaction, people often rely on expectations about the traits of other people based on markers of social group membership, i.e., stereotypes. Although the influence of stereotypes on social behavior is widespread, key questions remain about how traits inferred from social group membership are instantiated in the brain and incorporated into neural computations that guide social behavior. Here, we show that the human lateral orbitofrontal cortex (OFC) represents the content of stereotypes about members of different social groups in the service of social decision-making. During fMRI scanning, participants decided how to distribute resources across themselves and members of a variety of social groups in a modified Dictator Game. Behaviorally, we replicated our recent finding that perceptions of others’ traits, captured by a two-dimensional framework of stereotype content (warmth and competence), biased participants’ monetary allocation choices in a context-dependent manner: recipients’ warmth increased advantageous inequity aversion and their competence increased disadvantageous inequity aversion. Neurally, representational similarity analysis (RSA) revealed that perceptions of others’ traits in the two-dimensional space were represented in the temporoparietal junction and superior temporal sulcus, two regions associated with mentalizing, and in the lateral OFC, known to represent latent environmental features during goal-directed outcome inference outside the social domain. Critically, only the latter predicted individual choices, suggesting that the effect of stereotypes on behavior is mediated by inference-based, domain-general decision-making processes in the OFC.

## Introduction

In daily human life, people frequently make decisions about how to treat other people. Whether these decisions are fleeting (e.g., “Do I hold open the door for the approaching person?”) or more consequential (“Whom should I hire?”), a hallmark of human social decision-making is flexibility: the ability to adapt our behavior to interactions with different individuals based on information about what those individuals are like. However, people’s assumptions about what others are like are not always accurate. In particular, they are known to be influenced disproportionately by cues to the person’s group membership, such as the person’s gender, age, nationality, or occupation (i.e., stereotypes)^1–4^, setting up the potential to perpetuate disparities in treatment across different social groups. Although an abundance of research in the behavioral sciences has examined when and how people stereotype others based on their group membership^5,6^ and documented treatment disparities in domains ranging from medicine to healthcare^7^, it has been a challenge to characterize the impact of stereotypes on social decision-making processes, including the computational mechanisms that mediate the influence of stereotype information on social behavior^8,9^.

A recent advance at the intersection of psychology and behavioral economics offers a new framework to test hypotheses about how stereotypes about others’ traits are incorporated into neural computations that guide contextually-flexible social behavior. This advance builds upon the observation that stereotypes are structured along core dimensions of trait perception, such as warmth (the degree to which people have good intentions toward others) and competence (the degree to which people are capable of acting on their intentions)^5,6^. Using a novel modeling approach, we recently characterized how trait perceptions interact with the decision context to guide people’s resource allocation behavior^10^ toward members of different social groups^8^. Specifically, by incorporating stereotypes about others’ warmth and competence into a computational model of social valuation^11,12^, we found that these two dimensions of stereotype content exerted dissociable, context-dependent effects on individuals’ aversion to different forms of inequity: people were averse to receiving more money than stereotypically warm others, and people were averse to receiving less money than stereotypically competent others. In turn, this approach made it possible to predict with high accuracy not only individuals’ behavior toward a wide variety of social groups in a laboratory setting but also people’s treatment of members of different social groups in labor and education settings^8^.

This evidence points to the possibility that assumptions about others’ traits may be represented in the brain in a way that (i) corresponds to a dimensional structure of stereotype content and (ii) enables stereotypes to exert influence on the computations underlying social decisions in a context-dependent manner. To test this, we used fMRI and representational similarity analysis along with a social decision task involving members of different social groups.

Our hypotheses build upon our recent behavioral findings along with previous neuroimaging research into trait perception and stereotyping, on the one hand, and into value-based decision-making, on the other. First, a consistent set of brain regions including the temporoparietal junction (TPJ), superior temporal sulcus (STS), and medial prefrontal cortex (MPFC) is activated when people think about the minds of others and is therefore sometimes referred to collectively as the mentalizing network^13– 20^. Activations of the mentalizing network have been observed across a wide range of social task paradigms, including those that require inference of others’ traits based on their group membership (i.e., stereotyping)^21–25^. However, it remains unclear whether and how these regions mediate the effect of stereotypes on social decision-making, in large part because past studies of stereotyping have primarily involved passive viewing or basic judgments about others, making empirical characterization of behavior inapplicable; have focused mostly on *how active* different brain regions are, rather than on multi-dimensional trait representations^26^; and have primarily involved judgments about a small number of social groups (e.g., males versus females^24,27^), rather than a set of targets spanning the space of trait perception^28^.

Second, value-based decision-making has long been associated with processes in a set of frontostriatal regions, including the ventral striatum, the ventromedial prefrontal cortex, and the orbitofrontal cortex (OFC)^28–33^. A particularly intriguing area is OFC, which is thought to guide flexible, goal-directed decisions by representing defining features of the task or environment, often not directly observable but inferred, that are critical for inferring or imagining future decision outcomes^29–35^. Accordingly, the OFC may play a critical role in social behavior by representing others’ traits in ways that are behaviorally relevant. If so, OFC processes could plausibly serve as a route through which trait representations inform inference-based evaluation of overall decision outcomes in social contexts, including how subjectively rewarding particular monetary allocations with particular recipients will be. This account has the potential to unify the seemingly independent effects observed in past studies of social decision-making, which have shown that choices in the lab and field are modulated by overt characteristics such as race^36^, gender^37^, and attractiveness^38,39^, by suggesting that they share a reliance on core, underlying representations of perceived trait content.

Here we report evidence that neural representations of perceived trait content systematically bias social decisions in a way that relies on domain-general mechanisms of value-based decision-making in OFC. To do this, we conducted an fMRI experiment in which participants made decisions about how to allocate money across themselves and individuals from a variety of different social groups. Extending our previous behavioral findings^8^, we find that recipients’ perceived warmth increases advantageous inequity aversion and perceived competence increases disadvantageous inequity aversion. At the neural level, RSA revealed that stereotypic trait content was represented along the warmth and competence dimensions in the TPJ and STS, key regions in the mentalizing network, and in the OFC, a key region for goal-directed decision-making. Critically, we found that the representation in the OFC, but not in the other regions, predicted individual participants’ contextually-sensitive monetary allocation decisions. This suggests that, while regions of the mentalizing network may be involved in inferences about others’ traits, the effects of those trait perceptions on social decisions are mediated by domain-general mechanisms of inference-based, goal-directed decision-making centered in the OFC.

## Results

### Experimental paradigm

Participants (*n* = 32) played an extended version of the Dictator game in an fMRI experiment. The participant played the role of Dictator and, on each trial, decided how to allocate money between themselves and a recipient. To experimentally manipulate the participant’s perception of the recipient’s traits across trials, we provided one piece of information about the recipient’s social group membership (e.g., their occupation or nationality). We selected 20 social groups to span a wide range of social perception along the trait dimensions of warmth and competence, and ratings of their warmth and competence were collected in an independent, online sample (Fig. 1a)^8^. We also collected social perception ratings from our fMRI participants after scanning and confirmed that they were highly consistent with the independent ratings (Fig. S1), demonstrating the robustness of our social perception measures.

**Fig 1.**
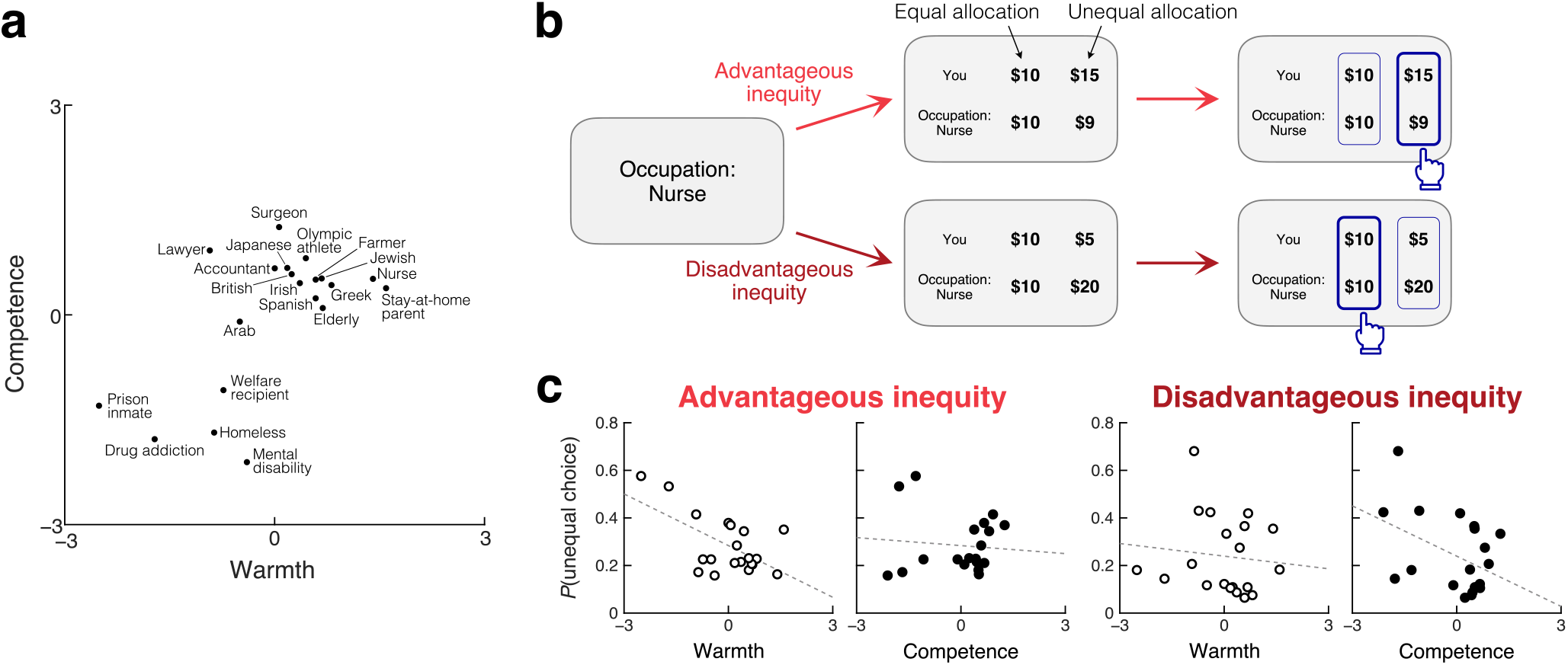
Experimental paradigm and behavioral results. **a**. Recipients in the Dictator game were identified by their social group membership. 20 social groups were chosen so that the recipient’s perceived warmth and competence were variable across trials. **b**. On each trial, the recipient’s social group was first presented, followed by two allocation options, one equal and one unequal. The participant was asked to make a binary choice. The unequal option allocated more money to the participant than the recipient in advantageous inequity trials (*top*) and less money in disadvantageous inequity trials (*bottom*). **c**. Participants’ allocation choices were influenced by the recipient’s perceived traits in a context-dependent manner. *Left*: In advantageous inequity trials, participants were less likely to choose the unequal option (and more likely to choose the equal option) when the recipient’s perceived warmth was higher (*r* = −.60, permutation *p* = .004), irrespective of their competence (*r* = −.09, *p* = .331). *Right*: in disadvantageous inequity trials, participants were less likely to choose the unequal option when the recipient’s perceived competence was higher (*r* = −.43, *p* = .040), irrespective of their warmth (*r* = −.11, *p* = .307).

On each trial, the participant was presented with the information about the recipient (e.g., “Occupation: Nurse”; “Nationality: Japanese”), and then with two monetary allocation options, between which they were asked to choose one (Fig. 1b). We manipulated these options so that we could empirically characterize the tradeoff between decision-making motives, i.e., maximization of one’s own payoff and concern for the inequity between oneself and the recipient. Specifically, in some trials, the participant chose between an equal allocation and an unequal allocation that created advantageous inequity (i.e., allocating more money to the participant than to the recipient); in other trials, the participant chose between an equal allocation and an unequal allocation that created disadvantageous inequity (i.e., allocating less money to the participant than to the recipient). This forced choice design allowed us to directly examine how participants’ preferences about advantageous and disadvantageous inequity depend on the recipient, and specifically, on the recipient’s perceived warmth and competence.

### Context-dependent effects of others’ traits on social decisions

Behaviorally, the recipients’ perceived warmth and competence exerted diverging effects on participants’ monetary allocation decisions; perceived warmth influenced choices in advantageous inequity trials, while perceived competence influenced choices in disadvantageous inequity trials (Fig. 1c). In advantageous inequity trials, participants were less likely to choose the unequal allocation (and more likely to choose the equal allocation) when the recipient’s perceived warmth was higher (Pearson’s *r* = −.60, permutation *p* = .004). Their choices about advantageous inequity were not correlated with perceived competence (*r* = −.09, *p* = .331), and the effect of warmth was stronger than that of competence (*p* = .004). Conversely, in disadvantageous inequity trials, participants were less likely to choose the unequal allocation when the recipient’s perceived competence was higher (*r* = −.43, *p* = .040). Their choices about disadvantageous inequity were not correlated with perceived warmth (*r* = −.11, *p* = .307), and the effect of competence was stronger than that of warmth (*p* = .049). Therefore, aversion to advantageous inequity increases with the recipient’s warmth, whereas aversion to disadvantageous inequity increases with the recipient’s competence. These behavioral results replicate our previous findings^8^ despite substantial differences in experimental design, including the use of binary forced choices between equal and unequal allocations (rather than continuous allocations) in the current study.

### Neural representations of others’ traits

Our behavioral findings show that perceptions of other people’s traits, guided by information about social groups and organized along distinct dimensions of warmth and competence, exert strong and dissociable effects on social decision-making processes as captured by our extended Dictator game. Accordingly, we next looked for neural representations of these perceived traits. To elucidate the representation of perceived traits and not payoff structures or decision processes, we focused on BOLD signals during the portion of each trial when the participant was presented with the recipient’s group membership, prior to the presentation of the allocation options (Fig. 1a). We looked for brain regions where two recipients that are similar to each other in perceived traits (e.g., an Accountant and a Japanese person, who are both perceived to have high competence and moderate warmth) evoke similar response patterns, and two recipient that are dissimilar in perceived traits (e.g., an Accountant and a Prison inmate) evoke dissimilar response patterns (representational similarity analysis; RSA^40^). We adopted a whole-brain searchlight approach that looked for brain regions where the representational dissimilarity matrix (RDM) of the local response patterns in a spherical searchlight was correlated with RDM of the perceived trait, defined by pairwise Euclidean distance in the two-dimensional space of warmth and competence (Fig. 2a). To construct the neural RDM, we quantified dissimilarity in response patterns using cross-validated Mahalanobis distance, which is a metric of the extent to which response patterns evoked by different recipients are consistently distinguishable across scanning runs^41^.

**Fig 2.**
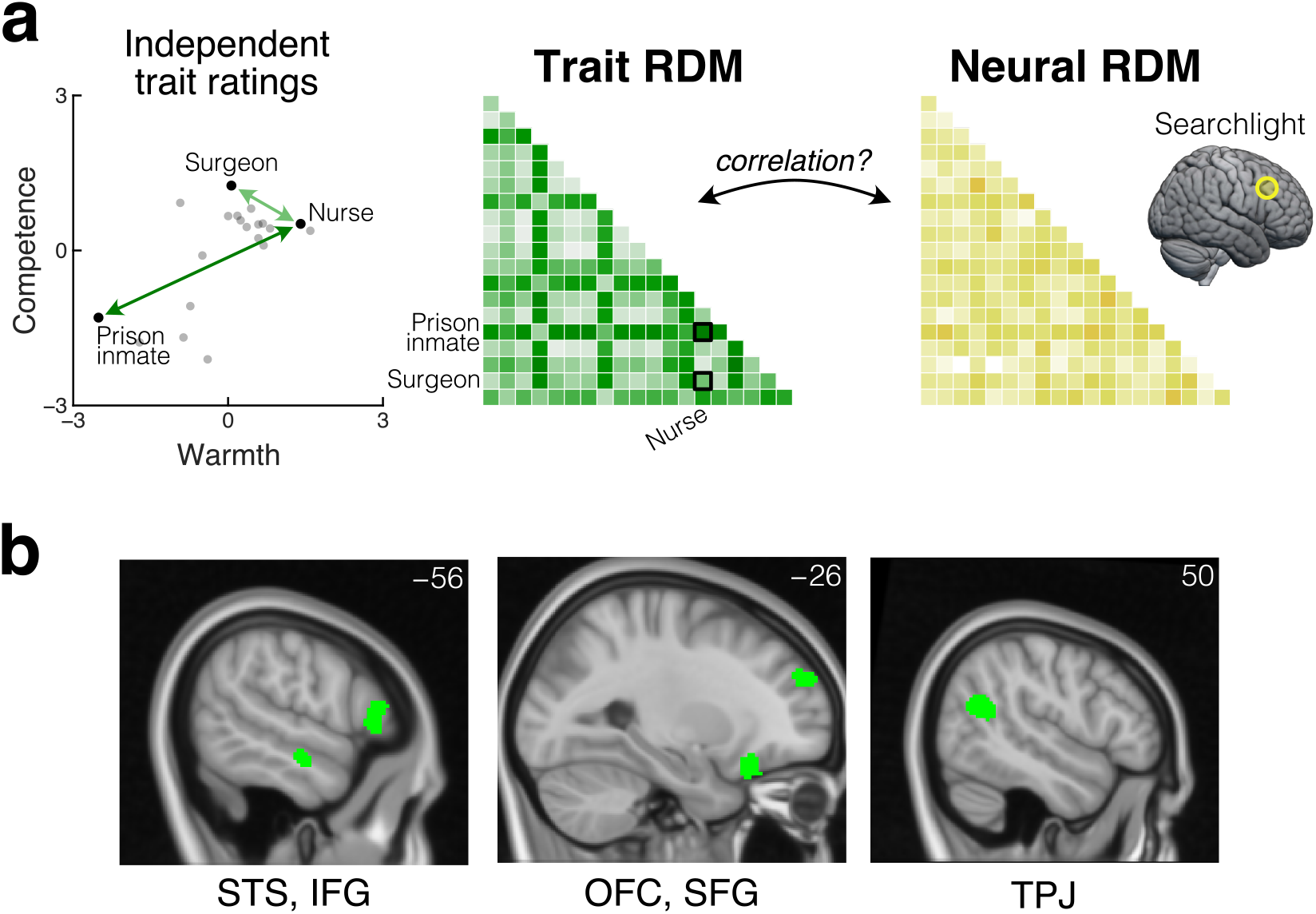
Neural representations of others’ traits. **a**. Whole-brain searchlight RSA looked for neural representations of the recipient’s perceived traits. The trait RDM was defined based on pairwise Euclidean distance in the two-dimensional space of warmth and competence. The neural RDM was computed for each searchlight based on pairwise cross-validated Mahalanobis distance between voxel-wise responses. **b**. Trait representation was found in left STS, left IFG, left OFG, left SFG, right TPJ, and right PMC (not shown) (whole-brain FWE-corrected TFCE *p* < .05).

Our RSA revealed that recipients’ perceived warmth and competence are represented in left lateral orbitofrontal cortex (OFC), which has long been associated with inference-based, goal-directed decision-making (threshold-free cluster enhancement [TFCE], whole-brain family-wise error [FWE] corrected *p* < .05). In addition to the OFC, perceived traits are also represented in several other regions, including those associated with mentalizing, such as the right temporoparietal junction (TPJ), left superior temporal sulcus (STS), left inferior frontal gyrus, left superior frontal gyrus, and right premotor cortex (Fig. 2b).

### Linking neural trait representations to choice behavior

Next, we investigated to what extent trait representations in these regions contributed to participants’ subsequent monetary allocation decisions (Fig. 3a). We reasoned that, if representations in any of the trait-representing regions (Fig. 2b) contribute to decision-making, then individual variations in local neural responses in such a region should predict individual variation in allocation choices. More specifically, if two recipients evoke similar response patterns in a particular region of a particular participant’s brain, and representations in that region contribute to decision-making in this context, then the participant should have treated those two recipients similarly. Likewise, recipients that evoke dissimilar response patterns in a given participant should have been treated dissimilarly by that participant. To test for such a relationship between neural responses and individual choices, we ran another RSA that examined the relationship between neural RDMs (on response patterns during the epoch of recipient identity presentation, as in the previous RSA) in each of the trait regions (Fig. 2b) and choice RDMs at the individual subject level (Fig. 3a). We visualized each participant’s choice frequency against each recipient (i.e., how often they chose the unequal allocation over the equal allocation) as a two-dimensional space, with choices in advantageous inequity trials on one axis and choices in disadvantageous inequity on the other axis. Pairwise Euclidean distance in this choice space was used to construct the individual choice RDM. To test the correlation between individual choice RDMs and neural RDMs above and beyond the population-level effects of warmth and competence, we obtained an FWE-corrected null-hypothesis distribution via permutation (randomly pairing choice and neural RDMs from different participants).

**Fig 3.**
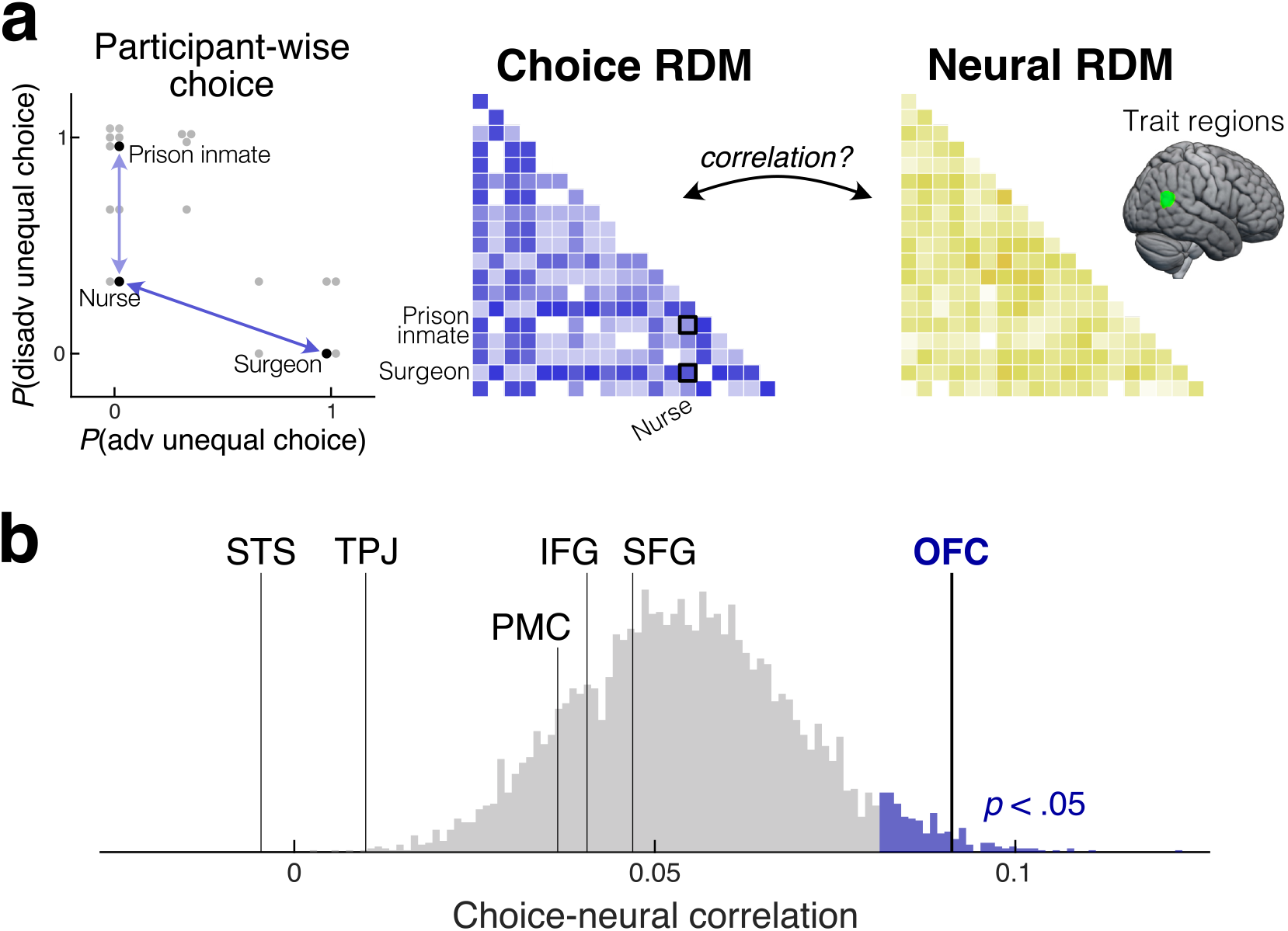
Correlation between neural representations of traits and individual choices. **a**. Relationship between individual-level allocation choices and response patterns in the regions that represent others’ traits (identified in **Fig. 2b**) was evaluated in the second RSA. The choice RDM was constructed for each participant based on pairwise Euclidean distance in the two-dimensional space of choice frequency in advantageous and disadvantageous inequity trials. Its relationship with the neural RDM in each trait region was measured by *Z*-transformed Spearman correlation. Shown is the data from one exemplar participant. **b**. The neural RDM in the OFC (*p* = .011), but not in any other region (*p* > .50), was significantly correlated with the individual-level choice RDM. Histogram: permutation-based FWE-corrected null hypothesis distribution.

This analysis revealed that only responses in the lateral OFC predicted individual allocation choices above chance (FWE corrected across the ROIs, *p* = .011; Fig. 3b). No other region exhibited a significant relationship with choices (*p* > .50). This suggests that the representation of the recipient’s traits in the lateral OFC contributes to the allocation decisions. Importantly, while our behavioral analysis revealed that the trait dimension (warmth or competence) that drives choices is *dependent on the decision context* (advantageous or disadvantageous inequity), responses in the lateral OFC were characterized by the two-dimensional spaces of traits (warmth and competence) and choices (advantageous and disadvantageous inequity), even before the participant was informed of the specific decision context. Taken together, these results suggest that the OFC plays a critical role in incorporating the perception of others’ traits into social decision-making in a highly flexible, goal-directed, context-dependent manner.

## Discussion

Adaptive social decision-making relies on information about others’ traits and mental states. However, we often need to interact with people with whom we have very little experience. In such cases, people sometimes rely on inferences derived from societally shared stereotypes based on cues to others’ social group membership^1–6,8^. Here, we identified a neural route through which stereotype content influences social decision-making. Using an extended Dictator game paradigm in which participants allocated monetary resources between themselves and various recipients identified by information about their social group membership, we first showed that people spontaneously treat others differently depending on their perceived traits in a context-dependent manner; advantageous inequity aversion increased with the recipient’s warmth, while disadvantageous inequity aversion increased with their competence. Using fMRI and RSA, we further showed that the recipients’ traits were represented in brain regions associated with both mentalizing (TPJ and STS) and goal-directed decision-making (OFC). Critically, the representation in the OFC was predictive of monetary allocation choices at the individual level. Using a permutation test, we confirmed that this relationship cannot be accounted for by population-level effects of warmth and competence, and instead implies that individual differences in the OFC signals are associated with those in decision-making. This shows that the OFC plays an important role in driving social decisions based on the perception of others’ traits.

Evidence that the lateral OFC mediates the effect of trait representations on social decision-making connects to a large body of evidence in humans and other species that the OFC contributes to goal-directed behavior. Goal-directed behavior is guided by inferred or imagined outcomes, as opposed to habitual behavior that is guided by cached values learned through trial and error. Previous studies used paradigms such as outcome devaluation or preconditioning to demonstrate that the OFC (in particular the lateral OFC) is necessary for goal-directed behavior in rats^42,43^, monkeys^44,45^, and humans^46–48^. Furthermore, recent neuroimaging and electrophysiological studies revealed that the OFC represents latent features of the environment, such as the hidden state of the current trial in sequential or learning tasks, that are not directly observable but are critical for outcome prediction^29,30,49–53^. Based on this evidence, a current influential hypothesis posits that the OFC represents aspects of the environment that are not fully observable but critical (or at least beneficial) for inference on future outcomes, and thereby guides flexible, goal-directed decision-making^31–35^.

Our findings, that the lateral OFC represents the perceived traits of others, and that this representation is predictive of individual choices regarding these others, are consistent with the hypothesized function of the OFC. First, recipients’ traits are not directly observable and instead inferred from information about their group membership. Second, decisions in the current paradigm are guided by inferences about how subjectively rewarding it would be to allocate money between the self and the recipient, as opposed to trial-and-error learning. Third, and most important, perceived traits affect inference-based evaluation of allocation outcomes, as demonstrated by the participants’ revealed preference in the current study as well as our previous studies with independent samples^8^. Taken together, this points to the possibility that the lateral OFC represent the recipient’s traits in the current experimental paradigm because they are critical variables for inference-based evaluation of resource allocations; it is likely that the OFC does not represent others’ traits in decision contexts that rely on other variables.

Other studies have also shown that the OFC is involved in incorporating perceptions of others’ traits into social decisions in a goal-directed manner. For instance, racial features of faces are represented in the OFC when participants chose whether to befriend them (goal-directed decision-making) but not when they judged whether they looked athletic (not goal-directed decision-making)^54^, and patients with lateral OFC damage are able to judge competence of faces but fail to incorporate it into voting decisions^55^. These findings, along with various social deficits exhibited by patients with OFC damage^35^, show that the role of OFC in inference-based, goal-directed decision-making extends to the social domain. Indeed, inference-based outcome evaluation is critical for a wide range of social decisions, since the social world is characterized by a high degree of uncertainty with complex latent structures (e.g., who are friends and who are foes) and countless unobservable variables (e.g., beliefs and preferences of individuals)^56,57^.

We also found neural representations of recipients’ traits in several regions outside the OFC. Among them, the right TPJ and the left STS are prominent areas in the mentalizing network, which is consistently activated when people infer others’ traits, including based on their group membership (i.e., stereotyping)^21–25^. Our results extend these previous findings by showing that multi-voxel response patterns in the TPJ and STS contain multi-dimensional information about the perceived traits of others. Interestingly, the STS (particularly its ventral bank, where we found trait representations) is anatomically connected to the lateral OFC in monkeys^58^, raising the possibility that the goal-directed representations in the OFC rely on inputs from the mentalizing network. In addition, the regions where we found trait representations outside the mentalizing network are also anatomically connected to the lateral OFC in monkeys^58–60^, and many of these regions are also functionally coupled with the lateral OFC in resting-state and task-based fMRI in humans^61,62^. Taken together, these findings suggest that the use of stereotypes in social decision-making relies on interaction between two key systems: one anchored on the mentalizing network, which is responsible for inferences about others’ traits, and the other primarily centered on the OFC, which incorporates the inferred traits into outcome inferences and evaluation in a context-dependent, goal-directed manner. This account is further supported by our finding that signals in the OFC, but not in other regions, are correlated with individual choices, which suggests that the OFC contributes to subsequent decision-making processes^63^.

Our findings open up a number of exciting questions for future research. First, future studies are needed to better understand the circuit-level mechanisms through which multi-dimensional representations in the OFC drive subsequent decision-making processes. For example, it is possible that the context-specific effects of social perception on behavior (warmth affects advantageous inequity aversion, while competence affects disadvantageous inequity aversion) could be mediated by flexible readout of the OFC signals by downstream regions^64^. Second, it remains an open question how trait representations in the mentalizing network and the OFC are constructed from semantic knowledge about social groups, possibly represented in the anterior temporal lobe^65–67^. Third, while we did not find evidence of trait representations in the hippocampus, a previous study reported that self-other relationships are represented in the hippocampus in a two-dimensional ego-centric space^68^. This raises the intriguing possibility that the OFC and hippocampus play complementary roles in social decision-making by representing the social world in different frames of reference^31,32,69–71^. Finally, our findings have the potential to inform future inquiry into the neuroscience of discrimination, for example by quantifying relationships between societal treatment of social groups and representations of their traits in the OFC^9,72,73^, as well as into disorders of social function, for example by separating social deficits arising from an atypical neural representation of others’ traits from those arising from an atypical integration of trait representations into value-based decision-making^74^.

Future research could also elucidate why trait representation was not observed in the MPFC in this context, at least at a standard statistical threshold for whole-brain analysis. Although the MPFC is also generally recruited during stereotyping^22–25^ and mentalizing^15–19,75,76^, it is possible that the MPFC contributes to stereotyping in a way that does not involve trait representations in a two-dimensional warmth-competence space^28,71,77,78^; that its contributions might be more specialized for inferences about individuals based on richer, more individuating information^79–82^; or that its involvement depends on the degree to which mentalizing is explicitly called for. For example, previous studies reported that the MPFC is more activated when participants receive explicit instructions to mentalize^83^, whereas the TPJ is consistently activated even when no explicit instructions or incentives for mentalizing are provided^75,84,85^. These possibilities further highlight the potential importance of goals and incentives in understanding the neural basis of social decision-making.

More broadly, while the current study focused on stereotypes, this is not the only route to trait inference. For instance, people often assume that others tend to hold attitudes or beliefs like their own (social projection), particularly when making inferences about individuals that are perceived to be similar to themselves^4,18,81,82,86^. Furthermore, for individuals with whom people interact extensively, trait information can be accumulated across learning from experience^65,87,88^. It remains an open question how trait information acquired through these different routes impacts social decisions at the cognitive and neural levels. For its part, the current study establishes how stereotypes drive social decisions via goal-directed representations in the OFC, forming the basis for a more comprehensive understanding of the neural mechanisms through which different types of social inferences affect social decisions across different contexts.

## Materials and Methods

All procedures were approved by the Institutional Review Boards at the University of California, Berkeley, and Virginia Tech.

### Participants

43 healthy people provided informed consent in accordance with the Declaration of Helsinki and participated in the experiment. Data from 1 participant were removed for image artifacts and data from an additional 10 participants were removed for excessive motion (showing frame-wise or cumulative displacement of >2mm in translation or >2.5 degrees in rotation), leaving data from 32 participants for analysis (22 female, 10 male, age: 18-64, mean = 27.5, standard deviation = 11.4).

### Task overview

Participants chose how to allocate monetary resources between themselves and a series of recipients in a modified dictator game. On each trial, the participant viewed one piece of social group information about the recipient for that trial (e.g., nurse, Japanese), along with two allocation options. In a majority of trials, one of the options provided an equal division of resources between the participant and the recipient, while the other option provided an unequal division of resources favoring either the participant (advantageous inequity) or the recipient (disadvantageous inequity). In the remaining trials, both options provided equal divisions in different amounts; these trials were only included to encourage the participant to pay attention to both sets of payoffs and were not included in the primary analyses in this paper (see Fig. S2c, d for behavioral data in these trials). In all cases, the participant decided unilaterally which option to choose, while the recipient had no ability to affect the outcome.

### Recipient identities

The recipient was described by one of 20 social group memberships, which were originally developed in our previous study^8^ to span a wide range of trait perceptions along the core dimensions of warmth and competence. The group membership was described by one of the following attributes: occupation (accountant, surgeon, lawyer, nurse, stay-at-home parent, Olympic athlete, farmer), nationality (Japanese, Irish, British, Spanish, Greek), ethnicity (Jewish, Arab), medical history (mental disability), age demographic (elderly), psychiatric history (drug addiction), housing status (homeless), financial status (welfare recipient), and legal status (prison inmate). The group membership was presented along with the attribute, e.g., “Occupation: Nurse” or “Nationality: Japanese”.

In all behavioral and fMRI analyses, we used ratings of these recipients’ warmth and competence collected from an independent sample in an online experiment (*n* = 252, Study 1b in our previous study^8^). To confirm that this independently measured social perception was shared by participants in the current fMRI experiment, we also asked these participants to rate recipients’ warmth and competence after the scan. We confirmed that the average ratings obtained in the current study were highly correlated with the independent ratings, demonstrating the robustness of our social perception measures (Fig. S1).

### Monetary allocation options

While the equal allocation option provided the same amount to the participant and the recipient ($10) across all trials, payoffs in the unequal allocation option were varied across trials. The payoff structure ([own payoff, the recipient’s payoff]) was either [$20, $5], [$15, $9], or [$14, $6] in advantageous inequity trials, and either [$5, $20], [$9, $15], or [$6, $14] in disadvantageous inequity trials. Therefore, in the advantageous inequity trials, the participant can maximize their own payoff by choosing the unequal allocation and maximize the recipient’s payoff by choosing the equal allocation. Conversely, in the disadvantageous inequity trials, they can maximize their own payoff by choosing the equal allocation and maximize the recipient’s payoff by choosing the unequal allocation.

### Procedure

Participants completed the task inside the MRI scanner and indicated their choices using a button box. The task was programmed in python using the Pygame package. Prior to scanning, participants were instructed that, although the monetary allocations in this task were hypothetical, they should indicate as honestly as possible which choice they would prefer if it were to affect the actual payoffs of themselves and the recipient. Throughout scanning, each of 8 payoff structures was presented once for each of the 20 recipients; in total, 8 × 20 = 160 trials were presented in a randomized order for each participant. The scanning consisted of two runs (80 trials each), with each recipient appearing four times per run.

In each trial, the participant was first presented with the recipient information (duration between 2.5 sec to 5.5 sec: varied across scanning runs and participants), and then with two allocation options, presented side by side. To mitigate cognitive load, the constant equal allocation [$10, $10] was always presented to the left, while the right option was varied across trials. After a delay (jittered between 3 sec and 6 sec), both options were outlined by blue boxes, which prompted the participant to indicate a choice by pressing one of two buttons. Participants were asked to press a button within 5 seconds; the trial was automatically terminated (and not repeated) when they did not press a button within that window.

### Behavioral data analysis

Economic theories of distributional preference posit that decision-making in the Dictator game is driven primarily by two factors: maximization of one’s own payoff and concern for the inequity between one’s own payoff and the recipient’s payoff^11,12^. They further posit that preferences regarding advantageous inequity are distinct from preferences regarding disadvantageous inequity^89,90^. In recent work, we found that aversion to advantageous inequity increases with the recipient’s perceived warmth (but does not depend on their perceived competence) and aversion to disadvantageous inequity increases with the recipient’s perceived competence (but does not depend on their perceived warmth)^8^. In that study, the participant decided how many tokens to share with the recipient in a continuous manner, and thus it was up to them whether and how often they created advantageous or disadvantageous inequity. We adopted a different task design in the current study, which used two-alternative forced choices regarding advantageous and disadvantageous inequity in separate trials, which allowed us to test the dissociable effects of perceived warmth and competence on inequity preference even more directly.

We counted how often the participants chose the unequal allocation over the equal allocation against each recipient in advantageous and disadvantageous inequity trials and tested their correlation with the perceived warmth and competence of the recipients for those choices (Fig. 1c). The statistical significance of the correlation was assessed via permutation (9,999 iterations). The same permutation test was also used to assess whether the effects of warmth and competence on choice frequencies were different from each other (i.e., statistical significance on the difference in correlations). While Fig. 1c shows choice frequencies marginalized over payoff structures in each trial type, the relationship with trait perceptions was robustly observed even when measured for each payoff structure separately (Fig. S2a, b).

### MRI data acquisition

MR images were acquired by a 3T Siemens Magnetom Trio scanner and a 12-channel head coil. A 3D high-resolution structural image was acquired using a T1-weighted magnetization-prepared rapid-acquisition gradient-echo (MPRAGE) pulse sequence (voxel size = 1 × 1 × 1 mm, matrix size = 190 × 239, 200 axial slices, TR = 2300 msec, TE = 2.98 msec). While participants completed the task, functional images were acquired using a T2*-weighted gradient echo-planar imaging (EPI) pulse sequence (voxel size = 3 × 3 × 3 mm, interslice gap = 0.15 mm, matrix size = 64 × 64, 32 oblique axial slices, TR = 2000 msec, TE = 30 msec). Slices were angled +30 degrees with respect to the anterior commissure-posterior commissure line to reduce signal dropout in the orbitofrontal cortex^91^.

### MRI data analysis: trait perception

We conducted a whole-brain searchlight Representational Similarity Analysis (RSA) to look for neural representations of the recipient’s perceived traits^40^. More specifically, we looked for brain regions in which voxel-wise local response patterns evoked by two recipients are similar (dissimilar) when their perceived traits are also similar (dissimilar) to each other. Our RSA formulated this relationship as the correlation between two representational dissimilarity matrices (RDMs), one that captures dissimilarity in trait perception (trait RDM) and one that captures dissimilarity in response patterns (neural RDM), in all possible pairs of recipients (20 recipients, 190 pairwise similarity measures).

For the trait RDM, pairwise dissimilarity in perceived traits was quantified as Euclidean distance in a two-dimensional space of perceived warmth and competence (Fig. 1a). Empirical measures of warmth and competence perceptions were originally obtained as numeric scores between 0 and 100^8^. We *z*-scored each dimension across the 20 recipients to construct the Euclidean space.

The neural RDM was computed at every voxel within grey matter in native space. Pairwise dissimilarity in voxel-wise response patterns was quantified as the cross-validated Mahalanobis (Crossnobis) distance in a gray-matter spherical searchlight (10mm radius). Crossnobis distance is an unbiased measure of the extent to which response patterns evoked by two recipients are *consistently distinguishable across scanning runs*^41^. We chose this distance measure over alternative measures because we were primarily interested in how recipients are *distinguished* in their neural representation, rather than how they are *similarly represented*. In our experiment, since each recipient was presented four times in each of the two scanning runs, we were able to cross-validate distance estimates across runs to mitigate spurious distance caused by noise (overfitting).

The pairwise Crossnobis distance was estimated following the formulae provided previously^41^. We first estimated voxel-wise response patterns evoked by each recipient in each scanning run using a GLM implemented in SPM12. To retain fine-grained signals as much as possible, minimal preprocessing (only motion correction) was applied to EPIs prior to the GLM. The GLM included the regressors of interest, modeling the presentation of each recipient using a box-car function that starts with the onset of the recipient presentation and ends with the onset of payoffs presentation, along with nuisance regressors modeling button presses. These regressors were convolved with the canonical double-gamma hemodynamic response function (HRF) and its temporal derivative. The GLM also included confound regressors for head motion (3 translations and 3 rotations, estimated in the motion correction procedure), 128-sec high-pass filtering, and AR(1) model of serial autocorrelation. The GLM coefficients of each recipient within the searchlight were then cross-validated across the two runs to obtain the Crossnobis distance. For Mahalanobis whitening, we estimated the covariance matrix in the searchlight using the GLM residuals and shrank it for invertibility^92^.

We computed Fisher-transformed Spearman correlation between the trait and neural RDMs at each gray-matter voxel. We discovered that the trait RDM inadvertently contained information about visual features of the recipient presentation on the screen, and specifically its character count. This visual confound was controlled by partialling out another RDM that captured the character count. The resultant correlation map was normalized to the standard MNI space based on the MPRAGE structural image of each participant and spatially smoothed (Gaussian kernel FWHM = 8 mm) using SPM12. For the population-level analysis, a cluster-level permutation test was conducted using FSL randomise (threshold-free cluster enhancement [TFCE], whole-brain FWE corrected *p* < .05, 4,999 iterations).

### MRI data analysis: correlation with individual choices

To look for evidence that any of the regions that represented the perceived traits (Fig. 2b) contributed to the subsequent monetary allocation decisions, we ran another RSA which tested the correlation between neural RDMs and choice RDMs. We predicted that, if a region contributed to the decisions, local response patterns evoked by two recipients in one participant’s brain would be similar (dissimilar) to each other when the participant treated them in a similar (dissimilar) manner in their allocation choices.

The individual choice RDM was built on the frequency at which each participant chose the advantageous or disadvantageous unequal allocation for each recipient. Pairwise Euclidean distance was measured in the two-dimensional space of the observed choice frequencies, one dimension for advantageous inequity trials and the other dimension for disadvantageous inequity trials. Since each recipient was presented in three advantageous inequity trials and three disadvantageous inequity trials, the choice frequency on each dimension was either 0, 1/3, 2/3, or 1.

These individual-level choice RDM were then correlated with neural RDMs in the regions identified by our first RSA as containing representations of others’ traits. Binary masks were functionally defined in standard MNI space based on the aforementioned population-level statistics (TFCE, whole-brain FWE corrected *p* < .05) and converted to the native space of each participant’s brain using SPM12. The *z*-transformed Spearman correlation between the choice and neural RDMs was averaged across all voxels in the native-space masks.

In order to test whether neural response patterns predicted individual choice patterns *above and beyond* the population-level effects of warmth and competence, we conducted a permutation test, randomly pairing choice and neural RDMs from different participants (4,999 iterations). To control for multiple comparisons across ROIs, the null-hypothesis distribution was constructed by taking the highest population average of correlation scores across the ROIs in each permutation iteration.

## Acknowledgments

The authors thank Nakyung Lee and Pierre Karashchuk for assistance with paradigm development; Duy Phan, Amanda Savarese, and Cassandra Carrin for assistance with data collection; and Dilara Berkay for helpful input and assistance with the preparation of fMRI data for analysis.

## Supplementary Figures

**Fig. S1.**
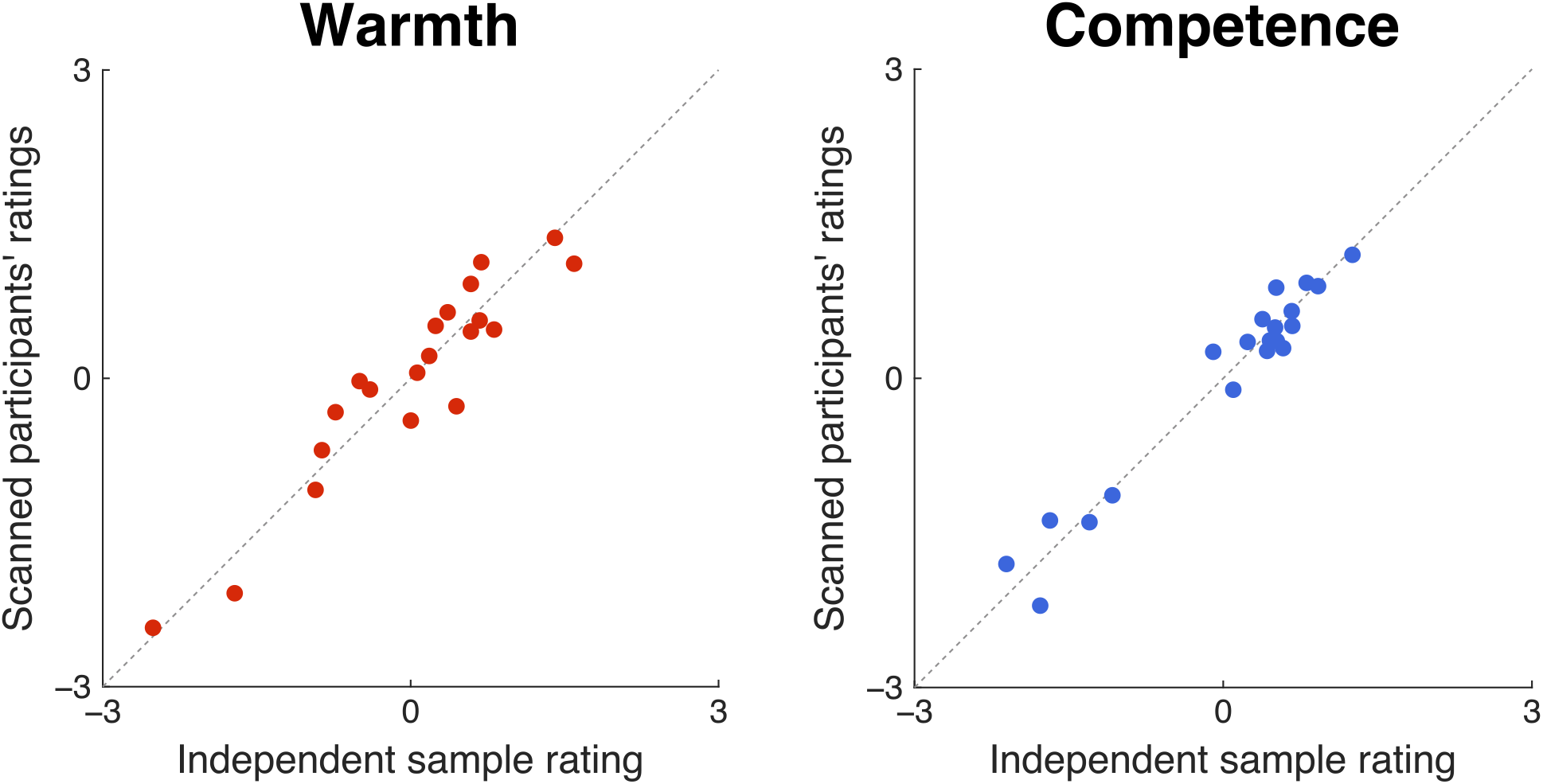
Consistency in trait perception. In all behavioral and fMRI analyses, we used ratings of warmth and competence from our previous study (Jenkins et al., 2018, Study 1b, *n* = 252; x axis). We also collected ratings from our participants after scanning (*n* = 32; y axis). These two sets of ratings are highly correlated (warmth: Pearson’s *r* = .943, competence: *r* = .978), demonstrating the robustness of trait perceptions.

**Fig. S2.**
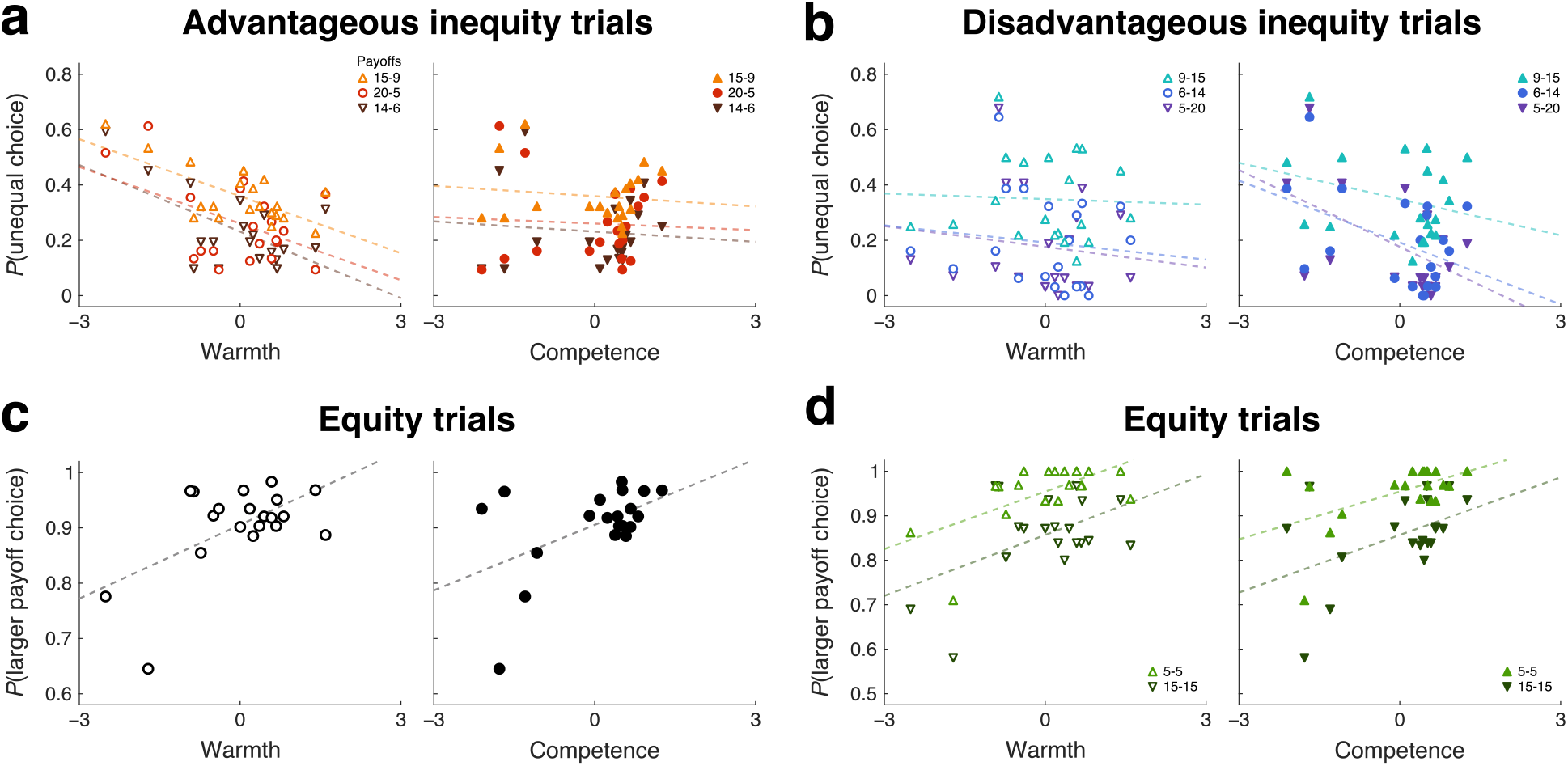
The effect of perceived traits on monetary allocation choices, separately for each payoff structure. **a**. In advantageous inequity trials, the unequal self-recipient allocations were either $15-$9, $20-$6, or $14-$6. Consistent patterns were observed across these payoff conditions; participants were less likely to choose the unequal allocation as the recipient’s perceived warmth was higher (*left*, 15-9: Pearson’s *r* = −.68, permutation *p* = .001, 20-5: *r* = −.47, *p* = .021, 14-6: *r* = −.60, *p* = .004) irrespective of the recipient’s perceived competence (*right*, 15-9: *r* = −.12, *p* = .285, 20-5: *r* = −.06, *p* = .386, 14-6: *r* = −.09, *p* = .331), and the effect of warmth was stronger than competence (15-9: *p* = .001, 20-5: *p* = .017, 14-6: *p* = .004). **b**. In disadvantageous inequity trials, the unequal self-recipient allocations were either $9-$15, $6-$14, or $5-$20. Consistent patterns were observed across these payoff conditions, except that the competence effect did not reach statistical significance in 9-15; participants were less likely to choose the unequal allocation as the recipient’s perceived competence was higher (*right*, 9-15: *r* = −.28, *p* = .125, 6-14: *r* = −.44, *p* = .036, 5-20: *r* = −.52, *p* = .018) irrespective of the recipient’s perceived warmth (*left*, 9-15: *r* = −.04, *p* = .417, 6-14: *r* = −.12, *p* = .287, 5-20: *r* = −.14, *p* = .265), and the effect of competence was stronger than warmth (9-15: *p* = .120, 6-14: *p* = .054, 5-20: *p* = .024). **c**. In some trials, the participant was presented with two equal allocations (one option was $10-$10, and the other option was either $5-$5 or $15-$15). These conditions were only included to encourage the participant to pay attention to both sets of payoffs and were not discussed in the main text. In these trials, participants chose the option with higher payoffs more often when the recipient’s warmth was higher (*r* = .57, *p* = .009), and also when their competence was higher (*r* = .51, *p* = .022). The effects of warmth and competence did not differ significantly (*p* = .362). These results demonstrate that participants incorporated the recipient’s warmth and competence into their choices in a highly context-dependent manner. **d**. Consistent behavioral patterns were observed across both payoff conditions in the equity trials; the larger payoff frequency increased with warmth (*right*, 5-5: *r* = .63, *p* = .006, 15-15: *r* = .49, *p* = .022) and competence (*left*, 5-5: *r* = .52, *p* = .020, 15-15: *r* = .46, *p* = .033), and their effects were comparable (5-5: *p* = .287, 15-15: *p* = .440).

## Notes

### Competing Interest Statement

The authors have declared no competing interest.

